# Cortistatin exerts an immunomodulatory and neuroprotective role in a preclinical model of ischemic stroke

**DOI:** 10.1101/2024.02.07.579281

**Authors:** J Castillo-González, L Buscemi, P Vargas-Rodríguez, I Serrano-Martínez, I Forte-Lago, M Price, P Hernández-Cortés, L Hirt, E González-Rey

## Abstract

Ischemic stroke is the result of a permanent or transient occlusion of a brain artery, leading to irreversible tissue injury and long-term sequelae. Despite ongoing advancements in revascularization techniques, stroke remains the second leading cause of death worldwide. A comprehensive understanding of the complex and interconnected mechanisms, along with the endogenous mediators that modulate stroke responses is essential for the development of effective interventions. Our study investigates cortistatin, a neuropeptide extensively distributed in the immune and central nervous systems, known for its immunomodulatory properties. With neuroinflammation and peripheral immune deregulation as key pathological features of brain ischemia, cortistatin emerges as a promising therapeutic candidate. To this aim, we evaluated its potential effect in a well-established middle cerebral artery occlusion (MCAO) preclinical stroke model. Our findings indicate that the peripheral administration of cortistatin at 24 hours post-stroke significantly reduces neurological damage and enhances recovery. Importantly, cortistatin-induced neuroprotection was multitargeted, as it modulated the glial reactivity and astrocytic scar formation, facilitated blood-brain barrier recovery, and regulated local and systemic immune dysfunction. Surprisingly, administration of cortistatin at immediate and early post-stroke time points proved to be not beneficial and even detrimental. These results emphasize the importance of understanding the spatio-temporal dynamics of stroke pathology to develop innovative therapeutic strategies. Premature interruption of certain neuroinflammatory processes might inadvertently compromise neuroprotective mechanisms. In summary, our study highlights cortistatin as a novel pleiotropic therapeutic approach against ischemic stroke, offering new treatment options for patients for whom early revascularization intervention is unsuccessful.

**Graphical abstract:** 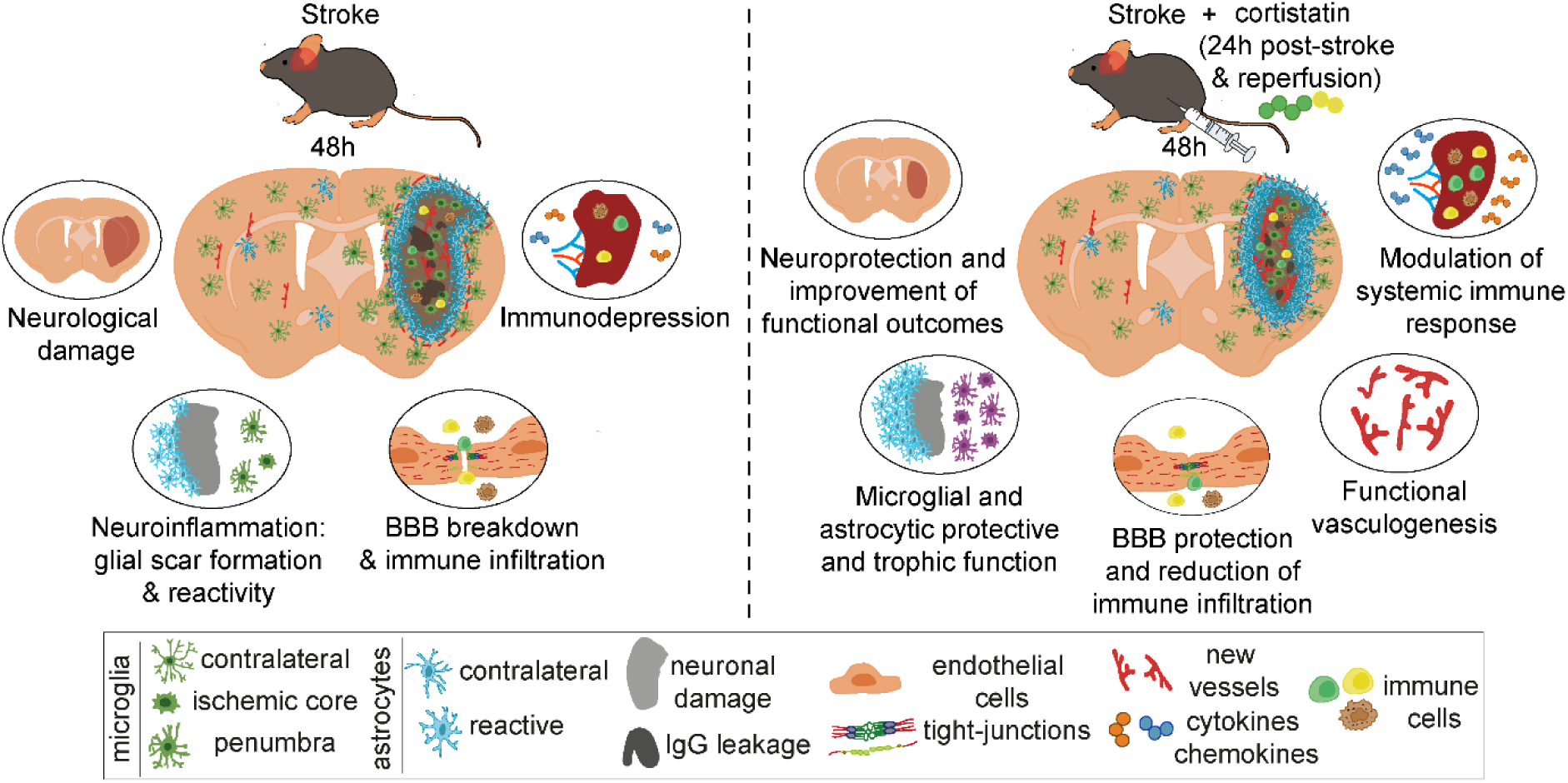

## 1. INTRODUCTION

Ischemic stroke results from a permanent or transient occlusion of a cerebral artery. The abrupt cessation of cerebral blood flow and subsequent reperfusion triggers a rapid and complex cascade of molecular and cellular events that may evolve over hours to days and weeks after the onset [1]. These pathophysiological events are diverse and include energy failure, oxidative stress, excitotoxicity, cell death, microglia reactivity, glial scar formation, blood-brain barrier (BBB) breakdown, and peripheral immune responses [2]. Notably, these processes are intricately interlinked, often unfolding concurrently within a brief time frame.

Despite significant efforts, stroke remains the second leading cause of death globally. Current treatments are limited to intravenous recombinant tissue plasminogen activator (rtPA) or endovascular thrombectomy. However, the narrow therapeutic window and rather limited efficacy sometimes restrict the utility of these treatments [3]. Furthermore, the intricate and still not fully understood spatiotemporal pathophysiology of stroke, with a plethora of underlying cellular and molecular processes involving the brain, vascular system, and peripheral organs, complicates its management. Consequently, there is an urgent need to thoroughly understand the neuroimmunological context and the factors involved at the different stages of ischemic pathology to develop effective therapeutic approaches.

Cortistatin (CST) is a cyclic neuropeptide broadly distributed in the central nervous system (CNS) and the immune system [4,5]. Despite sharing considerable homology, functions, and receptors (Sst1-5) with somatostatin, cortistatin exhibits unique functions in the nervous system and peripheral tissues [6,7]. These particular functions are mediated through interactions with receptors distinct from those of somatostatin, including the ghrelin receptor, the Mas gene-related receptor X-2, and a yet unidentified cortistatin receptor [5]. Notably, cortistatin displays potent anti-inflammatory properties, protecting against the development and progression of several inflammatory and autoimmune disorders (reviewed in [5]). Additionally, cortistatin treatment demonstrates immunomodulatory and neuroprotective effects in models of neuroinflammatory and neuroimmune disorders, such as multiple sclerosis [8], meningoencephalitis [9], excitotoxicity [10], neuropathic pain [11], and Parkinson’s disease [12]. In these conditions, cortistatin consistently reduces pro-inflammatory mediators, modulates glial cell responses, and maintains tissue-surveillance activities. Importantly, the absence of cortistatin in mice is associated with an exacerbated pro-inflammatory basal state in the CNS and periphery, along with decreased expression of neurotrophic factors [5,8]. Recently, our group has demonstrated a key role for cortistatin in maintaining BBB integrity, mediating anti-inflammatory endothelial functions, and facilitating BBB repair by stabilizing tight-junction proteins (TJs) and mitigating exaggerated immune responses after ischemic-like *in vitro* conditions [13]. However, no data has been reported about the influence of cortistatin on *in vivo* ischemic damage.

Therefore, given its neuroprotective, anti-inflammatory, and immunomodulatory properties, as well as its ability to regulate the BBB, we hypothesize that cortistatin could be a promising therapeutic agent affecting the intricate glial, vascular and peripheral responses underlying stroke pathophysiology.

## 2. MATERIALS AND METHODS

### 2.1. Peptides

Mouse CST-29 was acquired from Bachem. The lyophilized peptide was dissolved in distilled water (final concentration 10^-4^ M).

### 2.2. Animals

The procedures described in this study were approved by the Animal Care and Use Board and the Ethical Committee of the Spanish National Research Council (Animal Care Unit Committee IPBLN-CSIC # protocol CEEA OCT/2017.EGR), and the Vaud Cantonal Veterinary Office (protocol number 2017×6a) (Switzerland). All experimental protocols adhered to the guidelines specified in Directive 2010/63/EU of the European Parliament and the corresponding Swiss regulations regarding animal protection for scientific purposes. Mice were housed in groups of five animals per cage under controlled environmental conditions (60-70% relative humidity, 22 ± 1°C), and maintained on a 12-hour light/dark cycle (lights on at 7:00 a.m.). Food and water were provided *ad libitum*.

### 2.3. Transient middle cerebral artery occlusion model

Cerebral ischemia was induced in 12-week-old male C57BL/6J mice using the transient middle cerebral artery occlusion (MCAO) model, as adapted from Buscemi et al. [14]. Briefly, mice were anesthetized with isoflurane (1.5-2% in 70% N_2_O/30% O_2_) using a facemask. For post-surgical analgesia buprenorphine (0.025 mg/kg) was subcutaneously administered. A heating pad was used to maintain body temperature throughout the surgery. Regional cerebral blood flow (CBF) was monitored using laser Doppler flowmetry (moorVMS-LDF1, Moor Instruments) by fixing a probe on the skull over the core area supplied by the middle cerebral artery (approximately 1mm posterior and 6mm lateral from the bregma). Transient focal ischemia was induced by introducing a silicon-coated filament (701756PK5Re, Doccol) through the left common carotid artery into the internal carotid artery until it occluded the middle cerebral artery. The filament was left in place for 20 minutes before being withdrawn to allow reperfusion. Surgeries were considered successful when blood dropped below 20% of the baseline during occlusion, and raised above 50% of the baseline after filament withdrawal. Post-surgery, mice were housed with *ad libitum* access to food and water, with each cage containing water-softened chow and recovery gel.

Cortistatin (144μg/kg) was administered intraperitoneally at 0h and 24h (immediate treatment), 4h and 24h (early treatment), or 24h (late treatment) post-reperfusion. Control mice received an equivalent volume of saline solution.

### 2.4. Neurological score and behavioural tests

Neurological assessments were conducted at 24h following MCAO and prior to sacrifice at 48h. Stroke severity was assessed on a scale from 0 (no signs) to 3 (death) determining motor, balance, reflex, and sensory functions using a compendium of scales previously published [15,16].

To evaluate motor function and ataxia, the rotarod test was performed using a rotating cylinder apparatus (Ugo Basile, Gemonio, Italy) with a gradual acceleration protocol. The cylinder accelerated from 4rpm to 40rpm over 216s and then maintained a constant speed of 40rpm until reaching a 300s time limit. To ensure reliable results and exclude motivational factors, such as early falls, a training phase was conducted [17]. Mice underwent three consecutive daily training trials (with a 10-minute gap between them) for two days before MCAO (training trials, baseline). The longest latency to fall from the rod in three trials was recorded for each mouse. Mice underwent daily test trials at 24h and 48h after MCAO. For the test trials, the longest latency to fall from the rod or to complete two consecutive full turns while gripping the rod within the 300 seconds time limit was recorded for each mouse.

Additionally, muscle coordination and endurance were assessed using the wire-hanging test. Animals were suspended on a custom-made apparatus consisting of a single wire stretched between two posts above a soft ground. Mice were trained 24h before surgery and scored based on their performance in scaping (touching and reaching either the right or left post) or falling at 24 hours and 48 hours after MCAO [15]. The time elapsed until they escaped or fell was recorded. The test concluded either after 180 seconds or when a mouse had fallen off the wire ten times.

### 2.5. Sacrifice and infarct volume measurement

Mice were euthanized by intraperitoneal injection overdose of ketamine (100mg/kg, Ritcher Pharma)/ xylazine (10mg/kg, Fatro Iberica) at 48h post-MCAO surgery, followed by intracardial perfusion with 4% paraformaldehyde (PFA). Brains were collected and post-fixed in 4% PFA for 24h at 4°C, and then cryoprotected in 30% sucrose for 48h at 4°C. Subsequently, brains were frozen in isopentane on dry ice at −25°C and stored at −80°C until further processing. Coronal sections of 20µm thickness were obtained using a cryostat.

Cresyl violet (CV)-stained sections were prepared by gradual hydration in ethanol, stained with 1mg/ml of CV-acetate in distilled water, gradual dehydration in ethanol, and clearing in xylene [15]. Sections were imaged under a Leica DM2000 Transmit stereomicroscope at 2.5x magnification. The infarct volume was determined by multiplying the sum of the infarcted areas on each section (measured using the ImageJ Fiji software) by the section spacing.

### 2.6. Immunofluorescence

For immunofluorescence labelling, 20µm free-floating sections were washed with PBS and blocked and permeabilized with PBS containing 1% BSA, 0.1% Triton X-100, and 5% goat serum for 60min at room temperature (RT). Following this, sections were incubated overnight (O/N) at 4°C with primary antibodies anti-MAP2 (1:1000, Sigma), anti-Iba1 (1:1000, Wako), anti-GFAP (1:5000, Dako), anti-CD31/PECAM-1 (1:300,

BD), anti-ZO-1 (1:100, Fisher), or APC-labelled anti-CD45 (1:200, BD) in PBS containing 1% BSA, 0.1% Triton X-100, and 5% goat serum. After extensive washing with PBS, sections were incubated with the corresponding secondary Alexa Fluor-conjugated antibodies (1:1000, Sigma) and DAPI in PBS containing 1% BSA, 0.1% Triton X-100, and 2% goat serum for 60min. Sections were mounted with Mowiol. For IgG staining, an Alexa Fluor 594-conjugated goat anti-mouse IgG antibody was utilized without a primary antibody. Images were captured using a LEICA DMi8 S Platform Life-cell microscope at 20x magnification and a confocal Leica TCS SP8 STED microscope at 63x magnification.

### 2.7. Morphometric characterization

To comprehensively characterize morphological alterations in microglia and astrocytes post-stroke, we employed different morphometric analyses. Images of selected microglia and astrocytes within specific brain regions (*i.e.,* periinfarct or core area) were randomly selected, ensuring no overlap between cells, and that the entire nucleus and branches were included. A total of 15-30 cells per mouse (n=15-30) from 3-6 mice (N=3-6) were analysed. Image processing was conducted using ImageJ Fiji software, following a systematic protocol adapted from others [12,18,19]. Images were converted to 8-bit grayscale, pre-processed to minimize background noise and enhance contrast, and binarized by applying a pre-established threshold. Extraneous pixels not corresponding to cells were eliminated using the *Remove outliers* and *Despeckle* tools, with manual editing performed as necessary to address significant background noise.

Following image pre-processing, the Skeleton and Fractal Analysis plugins of ImageJ Fiji software [20] were used. Specific parameters assessed included the number of junctions (Skeleton Analysis) and the convex hull area (*i.e.,* the area containing the entire cell shape enclosed by the smallest convex polygon), fractal dimension (*i.e.,* index of complexity), span ratio (*i.e.,* the ratio of the axes of the convex hull from major to minor), and circularity (*i.e.,* how closely the shape of an object resembles a circle) (Fractal Analysis). For detailed procedures, refer to the user manual of the respective plugins [20].

### 2.8. Determination of the peripheral immune response

To evaluate the peripheral immune response, we determined the levels of systemic immune mediators and the spleen cellularity. First, to quantify the levels of peripheral inflammatory factors, blood samples were collected by cardiac puncture, allowed to clot for 2h at RT, and then centrifuged at 2,000rpm for 10min. The concentration of inflammatory mediators in the serum was determined by sandwich ELISA assay. Briefly, maxisorb plates (Millipore) were coated O/N at 4°C with the capture antibodies (anti-IL-6, anti-IL-12 from BD Pharmigen, or anti-MCP-1 from PeproTech). The plates were then washed with PBS containing 0.05% Tween20 and blocked with PBS containing 10% FBS for 2h. The samples and the recombinant standards were added to the plates and incubated O/N at 4°C. Following incubation, biotinylated detection antibodies (Serotec) were added for 1h at RT. After washing, plates were incubated with avidin-HRP (Sigma) (1:500 in blocking buffer) for 1h and then developed with ABTS substrate (Sigma). Absorbance was measured at 405nm using a spectrophotometer (Molecular Devices) and concentrations of inflammatory factors were determined based on the standard curve generated simultaneously.

Next, spleens were collected and weighed. Spleen tissue was gently ground with the frosted part of two slides onto a 100mm culture dish. The homogenized tissue was then transferred to RPMI medium supplemented with 10% FBS, 2mM L-glutamine, 1% P/S, and 50µM 2-mercaptoethanol. After allowing the suspension to sit for 5min on ice, the cell suspension was carefully collected and centrifuged at 1,670rpm for 10min at 4°C. To the resulting pellet, Ack buffer (150mM NH_4_Cl, 10mM KHCO_3_, 0.1mM Na_2_EDTA) was subsequently added for 10min at RT to lyse red blood cells. Next, RPMI medium was added, and cells were centrifuged at 1,670rpm for 10min at RT and counted.

### 2.9. RNA extraction and evaluation of gene expression

Following MCAO, brains were collected and divided into the contralateral and ipsilateral sides in both control and cortistatin-treated mice. For RNA isolation, brain tissues were placed in TriPure reagent (Roche). Then, cDNA synthesis was carried out using the RevertAid First Strand cDNA Synthesis Kit (ThermoFisher) with random hexamers in a CFX Connect QPCR System (Biorad) under the following conditions: incubation at 25°C/5 min, reverse transcription at 42°C/60 min, inactivation at 70°C/5 min. Gene expression analysis was conducted using conventional qPCR. SensiFast Sybr No-Rox kit 1x (Bioscience) was added to the cDNA corresponding to 20ng of RNA, along with the corresponding reverse and forward primers for each gene, and nuclease-free water to a final volume of 20µl. The CFX96 thermocycler (BioRad) was used to conduct the reaction under the following conditions: 94°C/5min (polymerase activation) followed by 40 cycles at 94°C/30s (denaturalization), 58°C/60°C/62°C/30s (primer annealing temperature depending on the gene), and 72°C/30s (extension). The comparative threshold cycle (Ct) method was performed for quantification. Each gene Ct values were normalized to those of the housekeeping gene *Rplp0* within each PCR reaction. Fold change was determined using the ΔΔCt method. The specific primers for each gene are provided in Additional file 1: Tables S1.

### 2.10. Statistical analysis

Data are presented as the mean ± SEM. All experiments were conducted in a blinded and randomized manner. The number of independent animals (N), and tissue sections and cell types (n) are indicated. Statistical differences between two groups were conducted using the unpaired Student’s t test for normal distribution and parametric data, or the non-parametric Mann-Whitney U-test. For comparing three or more groups that display normal distribution and parametric data, one-way ANOVA with Fisheŕs LSD test *post-hoc* tests for multiple comparisons was conducted. Three or more groups that display non-parametric data were evaluated by Kruskal-Wallis test with an uncorrected Dunn *post-hoc* test. Brown-Forsythe and Welch ANOVA with unpaired t-test and Welch’s correction *post-hoc* test were performed in the case standard deviations were not assumed to be equal. All statistical analyses were performed using GraphPad Prism v8.3.0 software, with P-values < 0.05 (one-tailed) considered statistically significant.

## 3. RESULTS

### 3.1. Identification of the optimal administration time of cortistatin in an acute stroke model

To investigate the potential therapeutic effect of the neuropeptide in ischemic stroke, we chose the MCAO model, a well-established preclinical model that closely mimics the human ischemic condition [21]. To consider the different phases of the pathology [2], we first administered cortistatin across different temporal contexts during stroke progression (Fig. 1A).

**Fig. 1.**
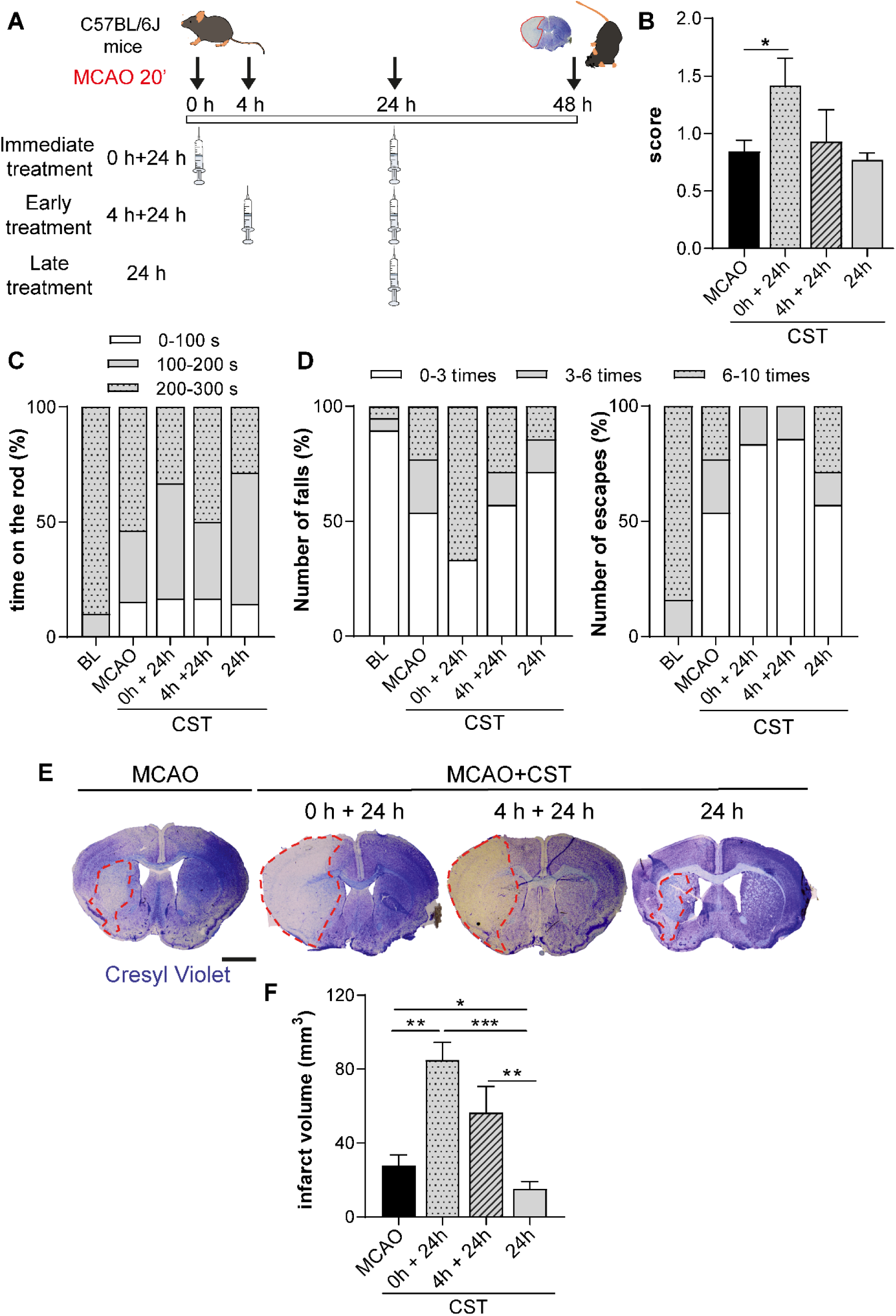
Optimization of the administration time of cortistatin in a preclinical stroke model. **A.** Schematic representation of the experimental timeline. C57BL/6J mice were subjected to MCAO and received either a saline solution (MCAO) or cortistatin (144μg/kg; MCAO+CST) at three distinct time intervals: immediately following reperfusion (0h + 24h), early after reperfusion (4h + 24h), and at late time post-reperfusion (24h). **B.** Neurological scores at 48h were assessed on a scale from 0 (no signs) to 3 (death). **C,D.** Behavioural performances were assessed by rotarod and wire-hanging tests, both at baseline (BL, pre-MCAO) and 48h post-MCAO. The time on the rod was evaluated in the rotarod test (**C**). In the wire-hanging test (**D**), the number of falls (**left**) and escapes (**right**) toward one of the posts were evaluated. Data are represented as the percentage of the total values based on the legend classification. N = 6-13 mice/group. **E.** Representative coronal brain sections from mice treated with either saline or CST at the specified times, stained with cresyl violet (CV). Ischemic lesions are outlined with dashed red lines. Scale bar: 1500µm **F.** Ischemic volume across the brain was determined by measuring ten CV-stained brain slices at 2-mm intervals per mouse. Data are expressed as mean ± SEM. \**vs.* MCAO-saline treated mice, \**p* ≤ 0.05, \*\**p* ≤ 0.01, \*\*\**p* ≤ 0.001. CST: cortistatin, MCAO: middle cerebral artery occlusion.

We administered the neuropeptide at three distinct time points: immediately post-MCAO (0h and 24h post-reperfusion), early post-MCAO (4h and 24h post-reperfusion), and late post-MCAO (24h post-reperfusion). The immediate treatment aimed to influence neuroinflammatory and neurodegenerative responses immediately after ischemia. The early treatment targeted pathological processes occurring within the first hours after hypoxia-reperfusion, while the late treatment focused on processes already established during the acute phase of stroke.

To determine the effect of cortistatin, we assessed neurological scores, behavioural impairments, and brain damage in C57BL/6J mice after administering cortistatin at the specified time points. First, neurological deficits resulting from stroke were evaluated within 24-48h after MCAO using a neurological scoring system encompassing motor, balance, reflex, and sensory functions graded from 0 to 3 (modified from [15]). As shown in Fig. 1B, MCAO control mice (saline-treated) exhibited a mean score of 0.5-1, indicating minor motor incoordination and slight hypomobility. Immediate cortistatin treatment significantly worsened neurological scores to 1-2, related to moderate intermittent circling, tremor twitches, and motor incoordination. However, early and late treatments did not significantly alter the neurological score compared to controls, despite the significant reduction in lesion size with late treatment (Fig. 1E,F).

We further evaluated motor function and muscle coordination using the rotarod and wire-hanging tests (Fig. 1C,D; Supplementary Fig. 1). In both tests, mice performance was measured 24h before the surgery (baseline, BL), and at 24 and 48h post-surgery.

As expected, both tests showed worsened performance in MCAO mice at 24h-48h post-surgery compared to BL (Fig. 1C,D, Supplementary Fig. 1). Specifically at 48h, saline-treated MCAO mice displayed a reduced time on the rod compared to BL (46.2% lasting less than 200 seconds *vs.* 90% of mice performing beyond this time point at BL), which was similar in the early treatment but reduced with the immediate cortistatin administration (50% and 71.4% of mice falling before 200 seconds, respectively) (Fig. 1C; Supplementary Fig. 1A). In the wire-hanging test, stroke control mice displayed a worsened performance and increased number of falls (only 53.8% falling less than 3 times *vs.* 89.5% of BL, and only 23.1% escaping more than 6 times *vs.* 84.2% of BL) (Fig. 1D, Supplementary Fig. 1B). Immediate treatment resulted in poorer performance compared to saline and other treatments (66.6% falling more than 6 times *vs.* 23.1% saline, and 83.3% escaping less than 3 times *vs.* 53.8% saline) (Fig. 1D, Supplementary Fig. 1B,C). Early treatment also showed similar trends in performance metrics (Fig. 1D, Supplementary Fig. 1C). However, although late treatment showed no beneficial effect on the rotarod test, it improved performance in the wire-hanging assay, with fewer animals falling more than 6 times compared to saline and other treatments (14.3% vs. 23.1% saline) (Fig. 1D, Supplementary Fig. 1B,C). Similarly, the percentage of animals escaping more than 6 times increased (28.6% vs. 23.1% saline or 0% for the other treatments) (Fig. 1D, Supplementary Fig. 1B,C).

Next, neurological lesions were quantified using CV staining, which differentiates intact neuronal cytoplasm (intense purple), from infarcted areas with dead neurons (faded purple) [22]. We observed that mice treated immediately after MCAO displayed larger infarct volumes when compared to MCAO controls (Fig. 1E,F). Remarkably, this treatment caused significant damage in the cortex and the striatum, spreading to the hippocampus area (data not shown), consistent with the observed worsened scores. Early treatment also led to larger lesions compared to MCAO-saline mice but to a lesser extent than the immediate treatment (Fig. 1E,F). In contrast, late treatment with cortistatin significantly decreased infarct volume, with only 23% of mice treated at 24h showing infarct volumes above 20mm³, compared to 44% of controls, 100% of immediately treated mice, and 66% of early-treated mice (Figure 1C,F).

### 3.2. Differential time administration of cortistatin affects stroke-derived glial responses

To investigate the underlying mechanisms by which cortistatin might differentially affect ischemic lesions and neurological deficits when administered at different times, we analysed the local neuroinflammatory response. In ischemic stroke, microglia are very early responders. These cells swiftly migrate to the infarct core, proliferating and adopting an amoeboid phenotype to engulf dead neurons [23]. Interestingly, immediate and early treatment with cortistatin resulted in reduced microglia density in the ischemic core compared to MCAO controls, while late treatment increased microglia density, potentially indicating a more physiological response to the injury (Fig. 2A,B).

**Fig. 2.**
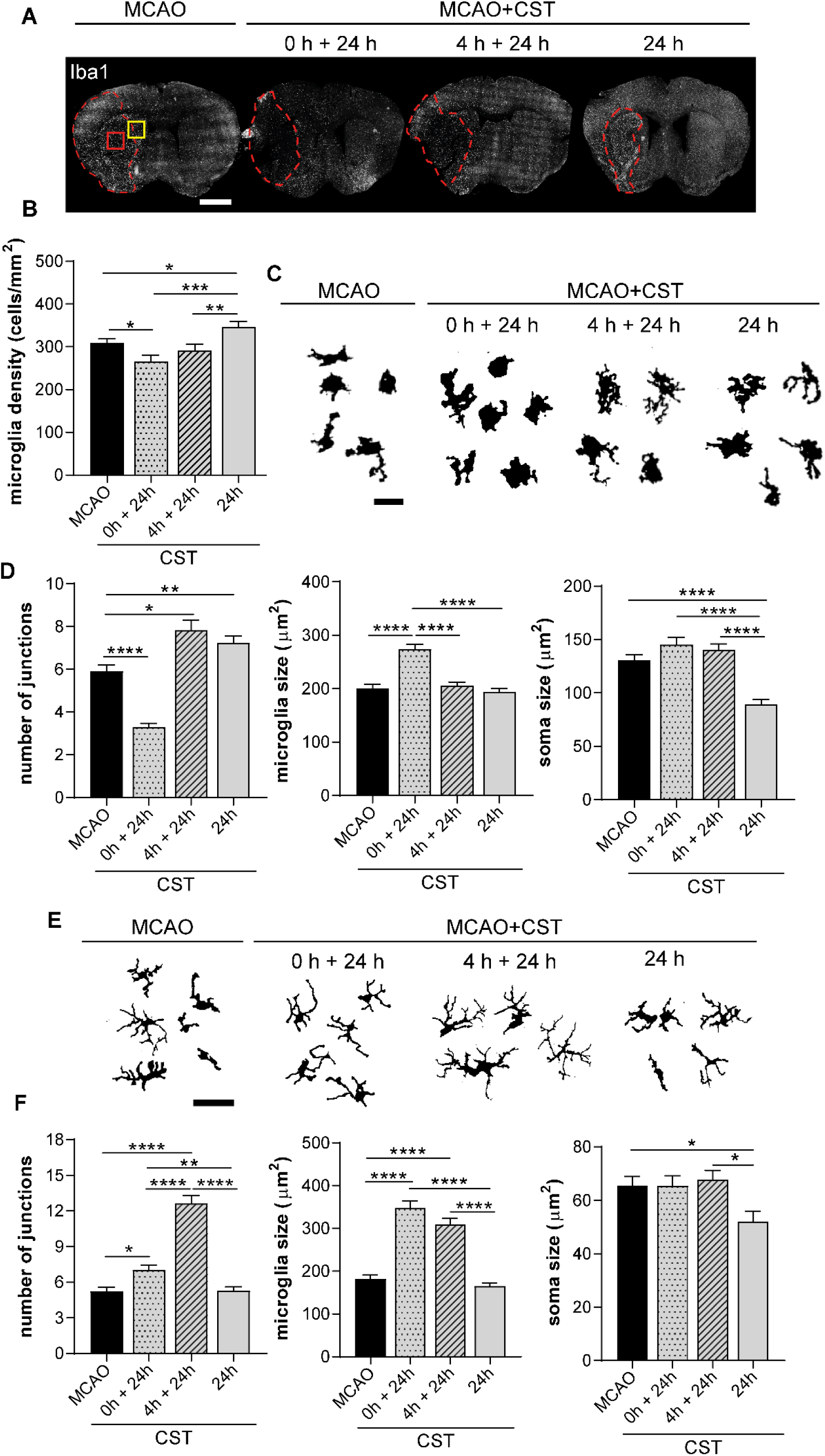
Time-dependent cortistatin treatment differentially regulates microglia response after stroke. **A.** Brain coronal sections of C57BL/6J mice were obtained after MCAO and subsequent treatment with either saline solution (MCAO) or cortistatin (144μg/kg; MCAO+CST) at three distinct time intervals: immediately following reperfusion (0h + 24h), early after reperfusion (4h + 24h), and at a late time post-reperfusion (24h). Microglia was identified by Iba1 staining. The ischemic core is represented by dashed red lines. Scale bar: 1,500μm. **B.** Density of microglia (cells/mm^2^) was measured in the ischemic core in the above-mentioned mice. **C-E**. Representative binary microglia from the different experimental groups from the ischemic core (red square) (**C**), or the penumbra (yellow square) (**E).** Scale bar: 30μm. Morphological parameters analysed (**D,F**). Skeleton Analysis was used to assess the number of junctions (**D,F** left panel). Cell area (**D,F** middle panel) and soma size (**D,F** right panel) were manually measured. n = 15-20 cells/mice; N = 3 mice/group. Data represent the mean ± SEM. \**vs.* saline \**p* ≤ 0.05, \*\**p* ≤ 0.01, \*\*\**p* ≤ 0.001, \*\*\*\**p* ≤ 0.0001. CST: cortistatin, Iba1: ionized calcium-binding adaptor molecule 1, MCAO: middle cerebral artery occlusion.

Since morphological alterations in microglia closely correlate with their reactivity states and serve as indicators of the severity of brain injury [18,24], we conducted Skeleton Analysis to evaluate microglia morphotypes. In the infarct core, microglia in saline-treated MCAO mice displayed a typical reactive amoeboid-like shape (Fig. 2C,D). Remarkably, immediate treatment with cortistatin led to a more pronounced reactive phenotype than controls, characterized by fewer junctions, and larger area. Conversely, early treatment resulted in a phenotype similar to MCAO mice, but with an increased number of ramifications, while late treatment was associated with an increased number of junctions and a significantly reduced soma, suggesting a transition to a less ameboid microglia state (Fig. 2C,D).

Microglia in the ischemic penumbra also undergo intricate morphological and functional changes [25]. We observed that immediate and early treatments increased the number of junctions and overall microglial size, indicating a hyper-ramified morphotype that was particularly pronounced in the early treatment. In contrast, late treatment resulted in smaller soma sizes, while the number of junctions and microglia area were similar to those in the MCAO control group, suggesting a different reactive state (Fig. 2E,F).

Parallel to the microglia response, astrocytes also become reactive and establish a glial scar surrounding the wound, containing the injury and preventing its expansion to adjacent still-viable regions [26] (Fig. 3A)

**Fig. 3.**
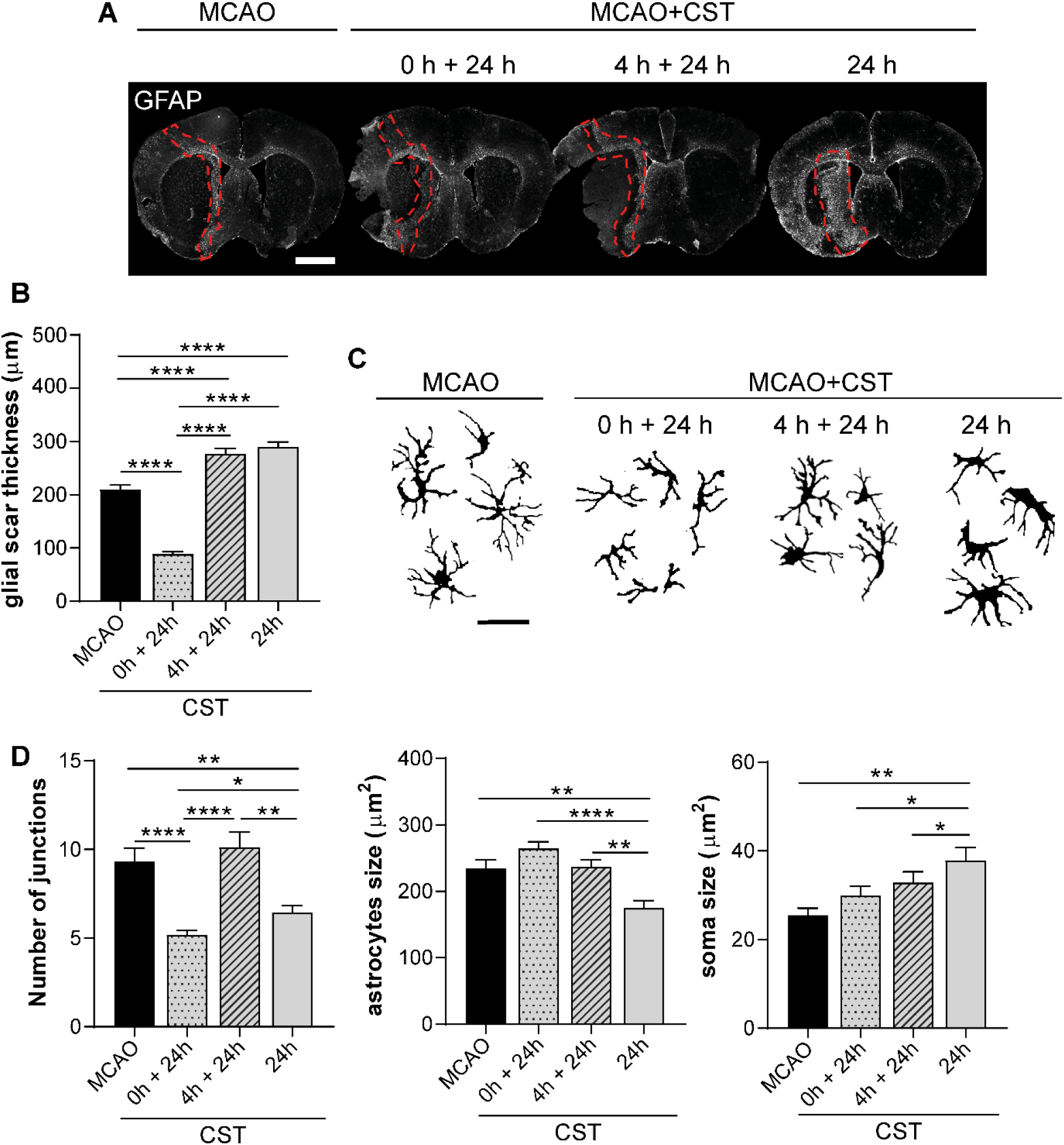
Time-dependent cortistatin treatment differentially impacts astroglial scar formation. **A.** Brain coronal sections of C57BL/6J mice were obtained after MCAO and subsequent treatment with either saline solution (MCAO) or cortistatin (115μg/kg; MCAO+CST) at three distinct time intervals: immediately following reperfusion (0h + 24h), early after reperfusion (4h + 24h), and at a late time post-reperfusion (24h). Astrocytes were identified by GFAP staining. Glial scar is outlined with dashed red lines. Scale bar: 1,500 μm. **B.** Glial scar thickness was quantified. N = 4-6 mice/group. **C.** Representative images of astrocytes in binary were extracted from the glial scar in the different experimental groups. Scale bar: 30μm. **D.** Evaluation of morphological parameters of astrocytes. Number of junctions (**D,** left panel) was evaluated by Skeleton Analysis. Cell (**D.** middle panel) and soma size (**D,** right panel) were manually measured. n = 15-20 cells/mice; N = 3 mice/group. Data are presented as the mean ± SEM. \**vs.* saline, \**p* ≤ 0.05, \*\**p* ≤ 0.01, \*\*\**p* ≤ 0.001, \*\*\*\**p* ≤ 0.0001. CST: cortistatin, GFAP: glial fibrillary acidic protein, MCAO: middle cerebral artery occlusion.

Remarkably, our study found that the immediate cortistatin treatment impaired glial scar formation, resulting in decreased scar thickness when compared to MCAO control mice. Nevertheless, early and especially late treatments significantly increased glial scar formation (Fig. 3A,B). Of note, the disorganized glial scar in the immediate treatment match with the severe neuronal lesion and the reduced microglial density in the core (Fig. 1E,F; 2B). We further analysed astrocyte morphology using Skeleton Analysis to explore the influence of cortistatin treatment in the astrocytic response. While astrocytes from MCAO mice displayed a high number of junctions, immediate treatment resulted in a significant reduction, suggesting a more quiescent astrocytic phenotype (Fig. 3C,D). Astrocytes from mice with early treatment were similar to the saline-treated MCAO mice, while late treatment showed astrocytes with reduced junctions and area, but increased soma size (Fig. 3C,D), suggesting a distinct reactive phenotype, which can be linked to the enhanced glial scar formation (Fig. 3A,B).

Collectively, these results suggest that during the hyperacute stroke phase, the initial glial response may be partially or fully deactivated by the potent anti-inflammatory effects of the neuropeptide when administered immediately or early. This deactivation leads to diminished protection of the damaged area, exacerbating neuronal lesions and motor deficits. In contrast, late cortistatin administration modulates the neuroinflammatory response differently, regulating glial cell hyperreactivity and/or proliferation, which results in reduced neuronal lesions and improved outcomes.

### 3.3. Treatment with cortistatin at later stages of ischemic stroke exerts neuroprotective effects

Based on these findings, we conducted a detailed investigation into the effects of cortistatin administered at 24h on neurodegeneration, glial population, BBB integrity, brain vasculature alterations, and local as well as peripheral immune responses at 48h following MCAO.

Our initial observations revealed that cortistatin-treated mice display reduced area of neuronal damage in both cortex and striatum compared to MCAO mice, as evidenced by MAP-2 neuronal staining (Fig. 4A,B), which was consistent with CV staining (Fig. 1E,F). Notably, ischemic injury coincided with decreased expression of neurotrophic factors such as BDNF and GDNF. Following treatment with cortistatin, the levels of these factors were significantly restored to baseline levels (Fig. 4C).

**Fig. 4.**
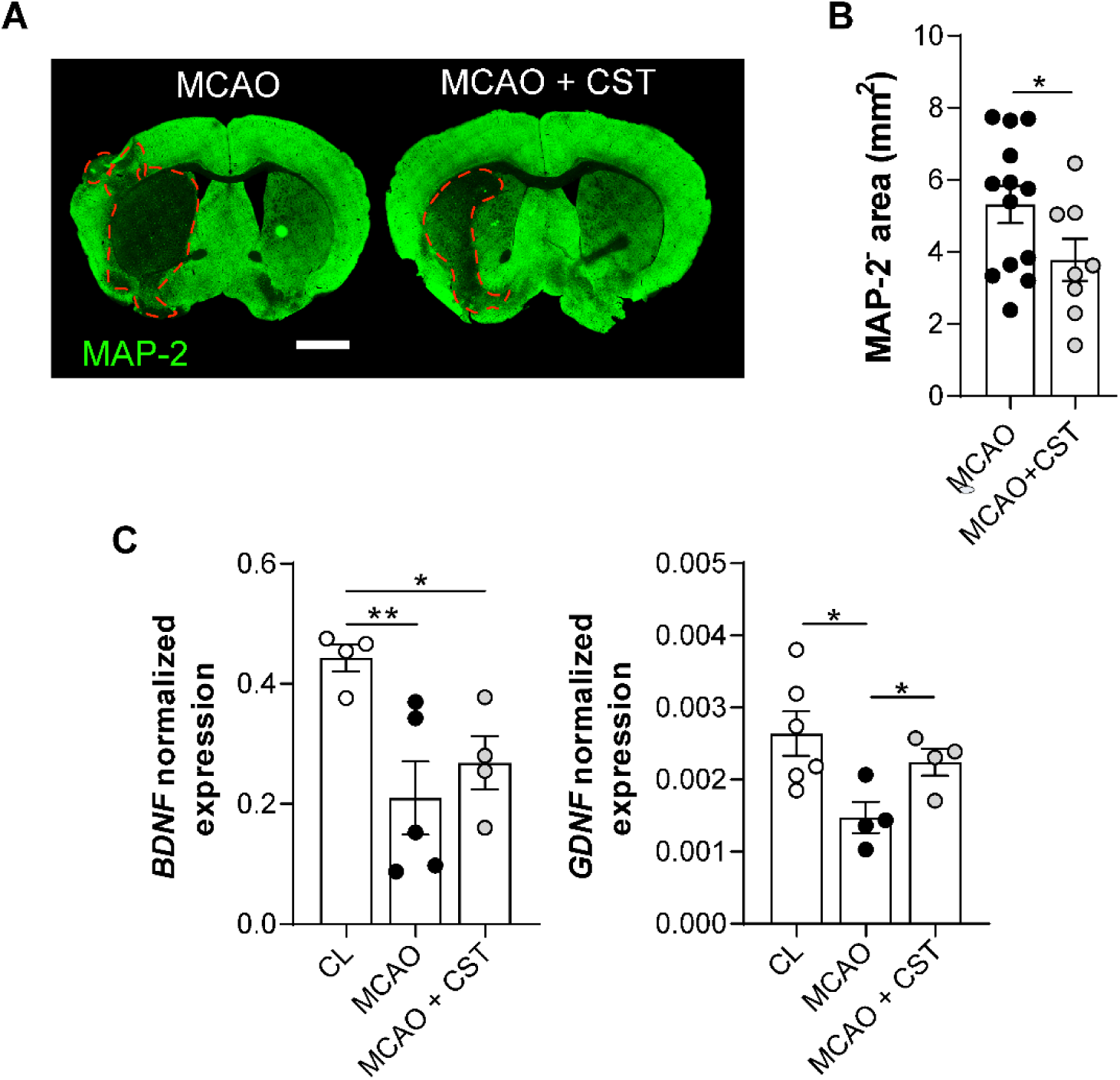
Late administration of cortistatin promotes neuroprotection. **A**. Brain coronal sections from C57BL/6J mice subjected to MCAO and treated with either saline (MCAO) or cortistatin (144μg/kg) (MCAO+CST) at 24 hours post-reperfusion and sacrificed at 48 hours. Sections were stained with MAP-2 to visualize neurons. Ischemic lesions, corresponding to a lack of MAP-2 staining, are delineated with dashed red lines. Scale bar: 1,500μm. **B**. Quantification of the ischemic lesion area. N = 13 (MCAO), 8 (MCAO+CST) mice/group. **C**. Gene expression levels in the contralateral (CL, uninjured) and ipsilateral (ischemic) brain hemispheres of mice treated with saline or cortistatin following MCAO were analysed by real-time qPCR and normalized to *Rplp0* expression. N = 4-6 mice/group. Data are presented as mean ± SEM with dots representing individual mice. \**vs*. saline, \**p* ≤ 0.05, \*\**p* ≤ 0.01. BDNF: brain-derived neurotrophic factor, CST: cortistatin, GDNF: glial cell line-derived neurotrophic factor, MAP-2: microtubule-associated protein 2, MCAO: middle cerebral artery occlusion.

### 3.4. Late treatment with cortistatin induces a neuroprotective microglial response in the acute stroke model

Next, we aimed to delve deeper into the role of cortistatin in modulating microglial morphotypes and functions, particularly focusing on its impact when administered at a later time post-stroke. To this aim, we complemented our initial morphometric analysis with a Fractal Analysis to evaluate various complexity parameters of microglia from the ischemic core and penumbra in MCAO mice treated with cortistatin (Fig.5; Supplementary Fig.2).

**Fig. 5.**
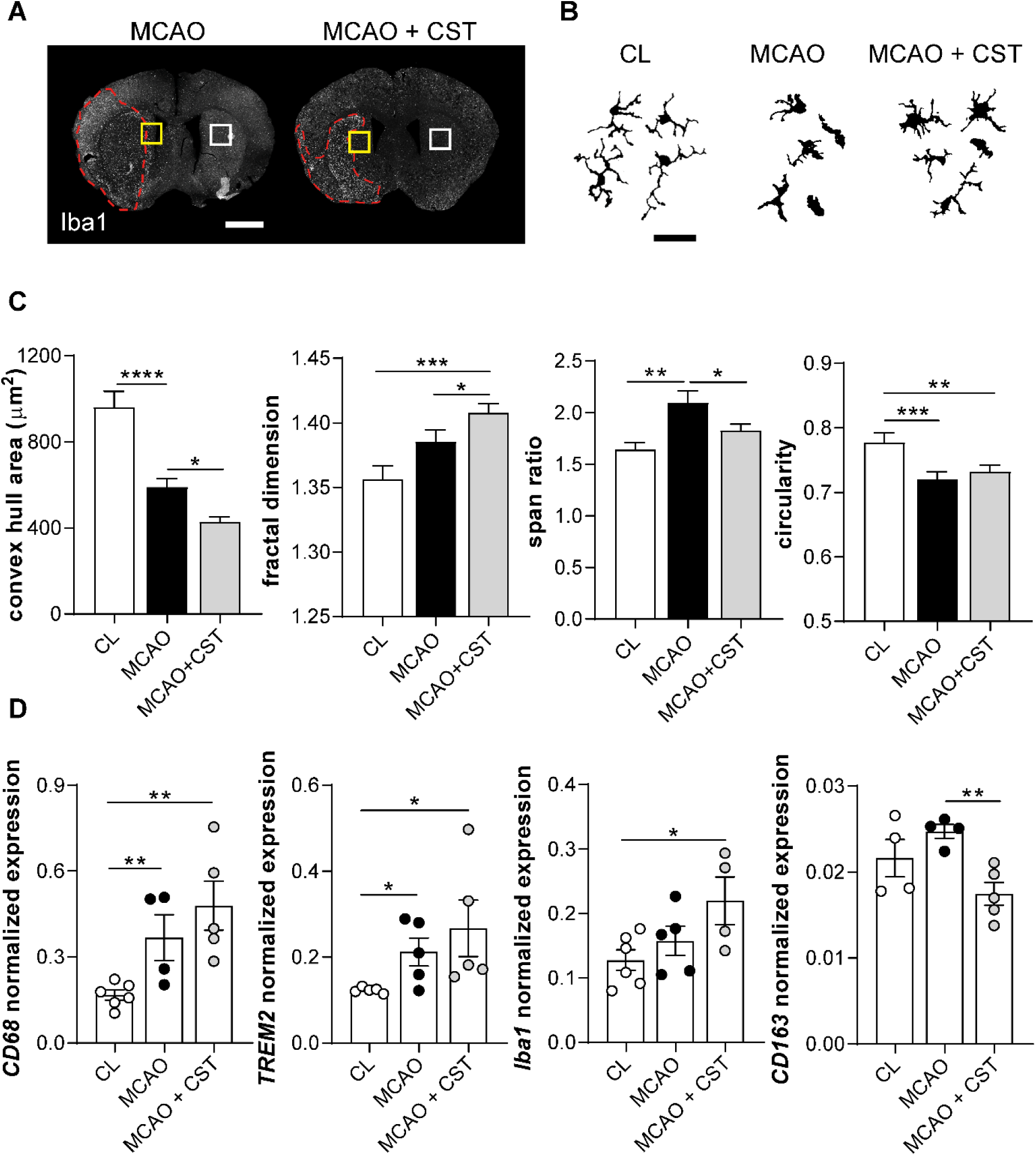
Late administration of cortistatin induces neuroprotective microglia. **A.** Brain coronal sections from C57BL/6J mice subjected to MCAO and treated with either saline (MCAO) or cortistatin (144μg/kg) (MCAO+CST) at 24 hours post-reperfusion and sacrificed at 48 hours. Microglia was identified by Iba1 staining. Ischemic core is represented by dashed red lines. Scale bar: 1,500 μm. **B.** Representative binary images of microglia from the ischemic penumbra (yellow square) and the contralateral side (CL) (white square) from the distinct experimental groups. Scale bar: 30 μm. **C.** Morphological parameters of microglia were analysed using Fractal Analysis including convex hull area, fractal dimension/complexity, span ratio, and circularity. n = 15-20 cells/mice; N = 3 mice/group. **D**. Gene expression levels of *CD68, TREM-2, Iba1,* and *CD163* in the contralateral and ipsilateral brain hemispheres of saline and CST-treated MCAO mice were measured by real-time qPCR and normalized to *Rplp0* expression (N=4-6 mice/group). Data are presented as mean ± SEM with dots representing individual mice. \**vs.* MCAO, \**p* ≤ 0.05, \*\**p* ≤ 0.01, \*\*\**p* ≤ 0.001, \*\*\*\**p* ≤ 0.0001. CD68: cluster of differentiation 68, CD163: cluster of differentiation 163, CST: cortistatin, Iba1: ionized calcium-binding adaptor molecule 1, MCAO: middle cerebral artery occlusion, TREM-2: triggering receptor expressed on myeloid cells 2.

MCAO animals exhibited a reactive microglial phenotype in the ischemic penumbra, characterized by a smaller territory area, enhanced complexity (as indicated by span ratio and fractal dimension), and decreased circularity compared to contralateral microglia (Fig. 5B,C). However, treatment with cortistatin seemed to mitigate this reactivity, evidenced by reductions in the soma size, junctions, territory area, and span ratio, along with an increase in complexity relative to saline-treated mice (Fig. 2E,F; Fig. 5C). These changes suggest a potential shift towards a less reactive phenotype or a different form of reactivity. In the ischemic core, although microglia from MCAO and cortistatin-treated mice looked similar, the treatment still led to distinct microglia with reduced soma size, increased junctions, and decreased span ratio (Fig. 2C,D; Supplementary Fig. 2A,B).

To further understand the functional implications of these microglial changes, we analysed the expression of different microglia markers in the ipsilateral hemisphere of saline and cortistatin-treated mice compared to the contralateral hemisphere (Fig. 5D). Following MCAO, there was enhanced expression of microglial reactivity markers such as *CD68* and *Trem2*. Remarkably, cortistatin treatment further upregulated the expression of *CD68*, *Trem2*, and *Iba1* compared to the contralateral hemisphere, and compared to MCAO mice (although the latter was not significant) (Fig. 5D). Interestingly, although no changes were observed for the gene expression of CD206 between healthy and ischemic brain samples (Supplementary Fig. 3A), there was a significant decrease in *CD163* expression in cortistatin-treated animals, advocating a potential promotion of both anti-inflammatory and increased phagocytic activity of these cells, consistent with the morphological findings and increased microglial density observed in the ischemic core (Fig. 2).

### 3.5. Late administration with cortistatin enhances glial scar formation and astrocyte reactivity

Given the collaborative role of microglia and astrocytes in the context of stroke to limit the damage and prevent its spread [26], we assessed the impact of late cortistatin treatment on the neuroinflammatory response of astrocytes (Fig. 6). Cortistatin treatment resulted in thicker glial scars with astrocytes exhibiting a reduced territory area, complexity index (fractal dimension), and circularity, and an increased span ratio, compared to control MCAO mice (Fig. 6A-C).

**Fig. 6.**
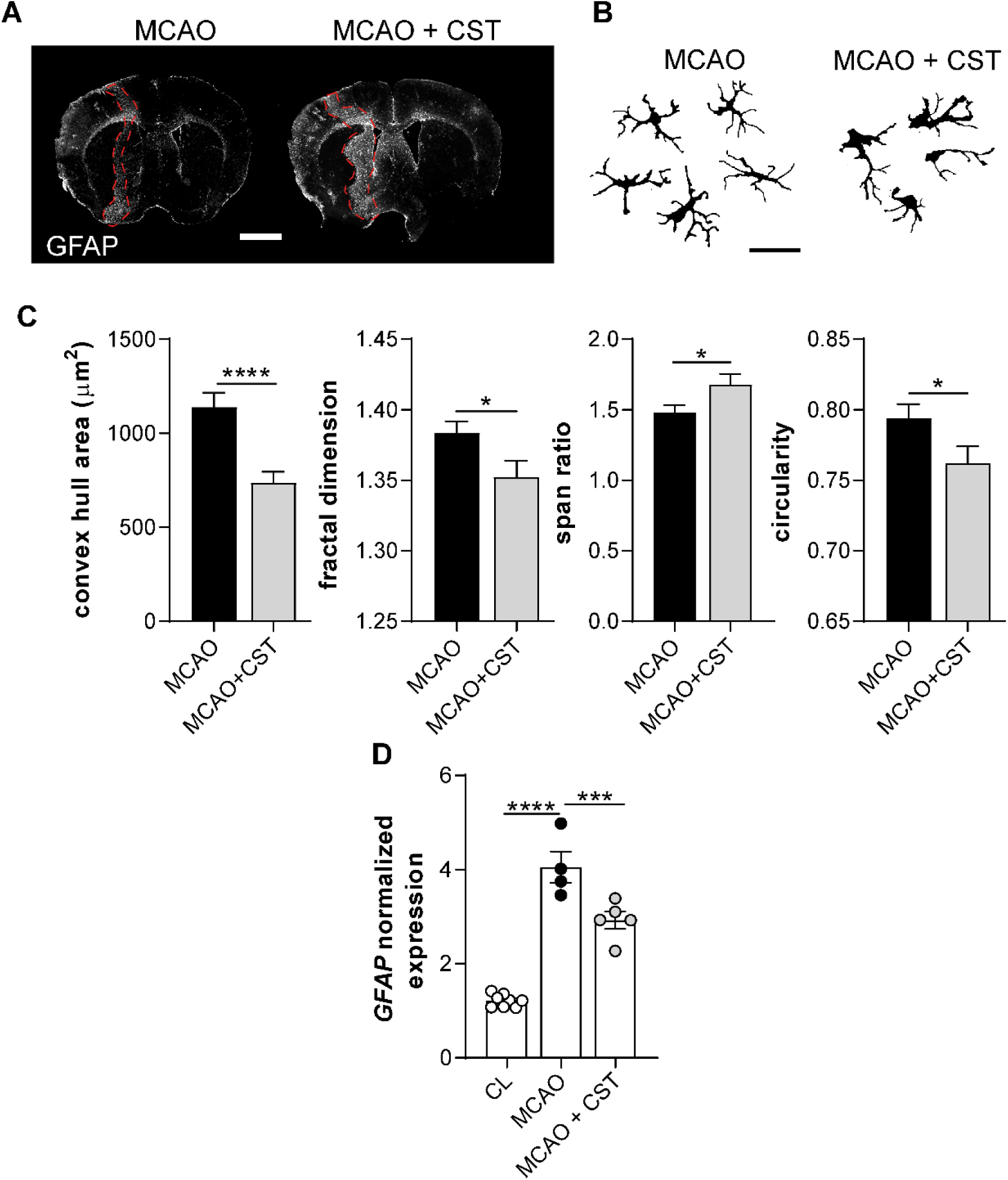
Late administration of cortistatin promotes protective astrocytic responses in ischemic stroke. **A.** Brain coronal sections from C57BL/6J mice subjected to MCAO and treated with either saline (MCAO) or cortistatin (144μg/kg) (MCAO+CST) at 24 hours post-reperfusion and sacrificed at 48 hours. Sections were stained with GFAP antibody to identify astrocytes. The astrocyte scar is indicated by dashed red lines. Scale bar: 1,500μm. **B.** Representative binary images of astrocytes from the glial scar were evaluated. Scale bar: 30 μm. **C.** Morphological parameters of astrocytes (convex hull area, fractal dimension/complexity, span ratio, and circularity) were quantified using Fractal Analysis. n = 15-20 cells/mice; N = 3 mice/group. **D**. Gene expression levels of *GFAP* in the contralateral and ipsilateral brain hemispheres of saline and CST-treated MCAO mice were quantified by real-time qPCR and normalized to *Rplp0* expression (N=4-6 mice/group). Data are presented as mean ± SEM with dots representing individual mice. \**vs* MCAO, \**p* ≤ 0.05, \*\*\**p* ≤ 0.001, \*\*\*\**p* ≤ 0.0001. CST: cortistatin, GFAP: glial fibrillary acidic protein, MCAO: middle cerebral artery occlusion.

Along with the fewer junctions, increased area, and enlarged soma shown in Fig.3, these parameters indicate that cortistatin induces a distinct morphotype of hypertrophic reactive astrocytes. To validate our findings, we examined the expression of *GFAP,* a well-known marker of astrocyte reactivity [27]. Following MCAO, both saline and cortistatin-treated mice showed increased *GFAP* expression compared to the contralateral side. However, cortistatin treatment significantly reduced *GFAP* expression relative to MCAO control mice. Thus, these results coupled with the upregulation of neurotrophic and glial factors after cortistatin treatment (Fig. 4,5), suggest that the reactive morphotype exhibited by these astrocytes may exert a role in the establishment of an efficient glial scar and potentially linked to reduced neuronal damage.

Together, these findings highlight cortistatin as a key regulator of glial responses in stroke pathology supporting neuroprotective properties.

### 3.6. Cortistatin protects against BBB breakdown and improves vascular dynamics after stroke

Stroke is accompanied by the breakdown of the BBB, contributing to neurovascular damage and complications. To thoroughly explore the extent of BBB dysfunction after ischemia in late cortistatin-treated mice, we evaluated IgG leakage. This is one of the key indicators of BBB breakdown, reflecting the extravasation of endogenous blood proteins. In our model, IgG leakage was prominently observed in the ipsilateral hemisphere after stroke, but significantly reduced following cortistatin treatment (Fig. 7A,B).

**Fig. 7.**
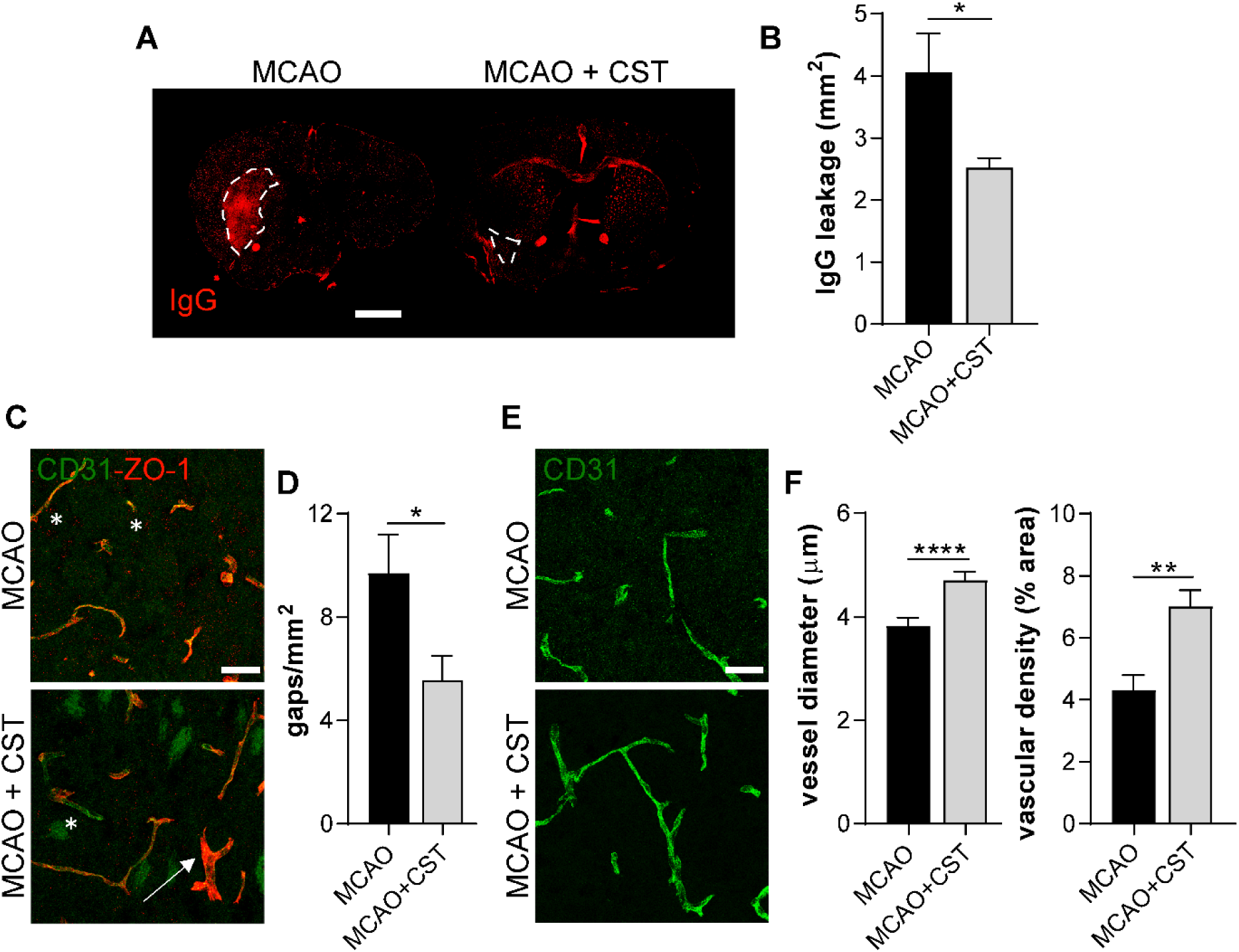
Cortistatin diminishes BBB disruption after stroke. **A.** Brain coronal sections from C57BL/6J mice subjected to MCAO and treated with either saline (MCAO) or cortistatin (144μg/kg) (MCAO+CST) at 24 hours post-reperfusion and sacrificed at 48 hours. Sections were stained with a fluorescently-labelled antibody against mouse IgGs. IgG leakage areas are outlined with dashed white lines. Scale bar: 1,500 μm. **B.** IgG^+^ area was measured as a quantification of IgG leakage from the blood. N = 4-8 mice/group. **C.** Representative images taken in the ipsilateral striatum of mice of the two experimental groups following MCAO and treated with CST, stained with CD31 antibody for brain endothelial cells (BECs, green) and ZO-1 (red) for TJs. ZO-1 gaps are highlighted with an asterisk, while ZO-1 coverage recovery is indicated with arrows. Scale bar: 30 μm. **D.** Quantification of the number of ZO-1 gaps per vessel area. n = 3 ROIs/section/mice; N = 3-5 mice/group. **E.** Representative images of BECs (CD31, green) from the ipsilateral striatum of MCAO or cortistatin-treated mice. Scale bar: 30 μm. **F.** Vessel diameter was manually measured, and vascular density was calculated as the % of positive CD31 staining of the total area of the image. n = 10-20 measures/vessel with 5-10 vessels/mice; N = 3-6 mice/group. Data are presented as the mean ± SEM. \**vs.* MCAO, \**p* ≤ 0.05, \*\**p* ≤ 0.01, \*\*\*\**p* ≤ 0.0001. CD31: cluster of differentiation 31, CST: cortistatin, IgG: immunoglobulin G, MCAO: middle cerebral artery occlusion, ZO-1: zonula-occludens 1.

After occlusion and reperfusion processes, hypoxic conditions particularly affect brain endothelial cells (BECs), disrupting TJs, leading to increased permeability and contributing to the development of vasogenic edema [28]. We observed that following MCAO, ZO-1 expression in the ischemic core showed disorganized patterns with increased gaps (Fig. 7C,D). Among TJs, disruption of ZO-1 has been extensively documented following stroke [29,30], playing a crucial role in BBB breakdown and subsequent recovery. Conversely, cortistatin treatment mitigated this disorganization, leading to a reduction in the number of ZO-1 gaps, thereby potentially improving TJ integrity (Fig. 7D). We also observed a decrease in the expression of *claudin-1* after MCAO *vs.* the contralateral side, with no alterations after cortistatin treatment (Supplementary Fig. 3B). Additionally, alterations in basement membrane proteins like collagen type I (Col1a2) were observed post-stroke, with cortistatin treatment showing a trend towards increasing *Col1a2* expression levels compared to MCAO alone (Supplementary Fig. 3B). In response to stroke, vascular dynamics undergo significant changes to promote angiogenesis and the modification of the vascular tree [31]. To investigate the importance of this process in stroke recovery, we examined the area occupied by vessels (CD31^+^) and their diameter in the striatal core (Fig. 7E).

Cortistatin treatment enhanced vessel diameter and vascular density in the ischemic striatum compared to saline-treated MCAO mice (Fig. 7F). This enhancement in vascular parameters may facilitate improved reperfusion to the damaged area, supporting post-stroke recovery mechanisms.

These findings suggest that cortistatin not only protects against BBB permeability breakdown but also promotes angiogenic and vascular remodelling events that aid in vascular repair and regeneration post-stroke.

### 3.7. Cortistatin modulates central and peripheral immune responses following stroke, reducing brain infiltration and immunodepression

After ischemia-reperfusion injury, the brain undergoes oxidative stress and neuronal death, triggering glial cell activation and the secretion of cytokines and other immune factors. Following MCAO, we observed increased expression of *TNF-α* and *IL-6* in the untreated ischemic brain hemisphere compared to the contralateral side, with no differences in *IL-10* expression. Interestingly, cortistatin treatment significantly elevated *TNF-α* and *IL-10* levels (Fig. 8A). Levels of *IL-1β* showed no significant differences between groups. Additionally, ischemia-induced arginase 1 (*Arg1*) expression, indicative of brain immune infiltration and stroke severity [32,33], was significantly downregulated by cortistatin (Supplementary Fig. 3C). This suggests that cortistatin influences the polarization of immune cells towards a neuroprotective phenotype.

**Fig. 8.**
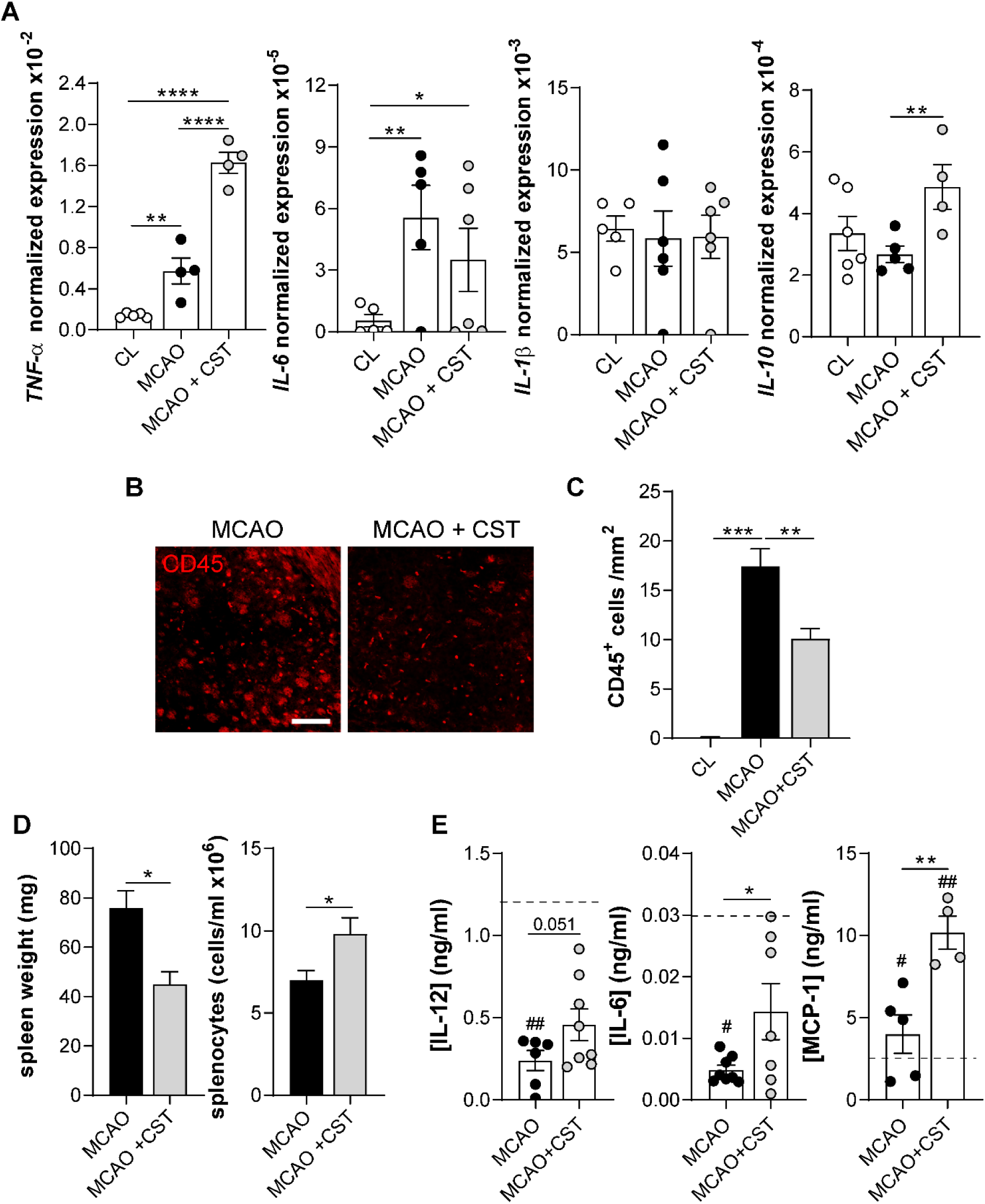
Cortistatin modulates immune infiltration and peripheral immune response after ischemic stroke. **A.** Gene expression levels for *TNF-α*, *IL-6*, *IL-1β*, and *IL-10* in the contralateral and ipsilateral brain hemispheres, of saline and CST-treated MCAO mice were quantified using real-time qPCR and normalized to *Rplp0* expression. N = 4-6 independent mice**. B.** Representative brain coronal sections stained with CD45 antibody (red) for leukocytes from C57BL/6J mice subjected to MCAO and treated with saline (MCAO) or cortistatin (MCAO+CST) at 24h post-reperfusion and sacrificed at 48h. Scale bar: 100 μm. **C.** Leukocyte density (number of cells/mm^2^) was measured in the striatum and the contralateral side (CL). **D.** Spleen weight and the number of splenocytes/ml were measured. N = 4-8 independent mice. **E.** Serum levels of IL-12, IL-6, and MCP-1 (ng/ml) in serum were measured using an ELISA assay. Dashed lines represent cytokine concentrations in healthy mice. N = 3-6 independent mice. Data are the mean ± SEM with dots representing individual mice. \**vs.* MCAO, ^#^*vs.* healthy mice *^,#^*p* ≤ 0.05, **^,##^*p* ≤ 0.01, \*\*\**p* ≤ 0.001, \*\*\*\**p* ≤ 0.0001. CD45: cluster of differentiation 45, CST: cortistatin, IL-6: interleukin 6, IL-1β: interleukin 1β, IL-10: interleukin 10, IL-12: interleukin 12, MCAO: middle cerebral artery occlusion, MCP-1: Monocyte Chemoattractant Protein-1, TNF-α: tumour necrosis factor-α.

Cytokines and DAMPs derived from brain damage can be released into the systemic circulation, getting to lymphoid organs and initiating the peripheral inflammatory response [34]. This triggers the quick adhesion of platelets and leukocytes to the endothelium facilitated by BBB disruption, enabling peripheral immune cells to infiltrate the brain [35]. To explore how cortistatin influences peripheral immune response dynamics, we measured cytokine levels in serum and assessed the infiltration of immune cells into the ischemic core. We observed that in MCAO mice, significant immune infiltration (CD45+ cells) occurred in the ischemic region, which was markedly reduced following cortistatin treatment (Fig. 8B,C). This suggests that cortistatin limits the infiltration of peripheral immune cells into the injured brain, possibly through its effects on BBB integrity.

The initial, short-lasting activation of systemic immunity by DAMPs and cytokines released in the bloodstream is swiftly followed by long-lasting immunodepression, characterized by lymphopenia and reduced functional activity of T cells and monocytes among others [36]. In our model, stroke-induced immunodepression was characterized by an oedematous and fibrotic spleen with decreased splenocyte numbers. Cortistatin treatment reversed these effects, significantly increasing the total number of splenocytes and reducing spleen weight after stroke (Fig. 8D). Furthermore, cortistatin treatment restored the reduced stroke-associated serum levels of IL-12 and IL-6 to homeostatic conditions, and increased MCP1 levels, indicating a systemic modulation of inflammatory responses (Fig. 8E).

In summary, cortistatin administered with an appropriate post-stroke timing exerts multifaceted beneficial effects. It enhances neuroprotective cytokine profiles in the brain, reduces neuronal damage, potentially through the modulation of microglia reactivity and enhancing the formation of the astrocytic scar, regulates BBB integrity preventing immune cell infiltration, and modulates systemic inflammatory responses. Additionally, cortistatin counteracts stroke-induced immunodepression by restoring splenic immune function. These findings underscore the therapeutic potential of cortistatin in stroke by not only protecting neuronal tissue and promoting vascular recovery but also by regulating immune responses to support overall recovery mechanisms.

## 4. DISCUSSION

The pathogenesis of stroke involves a complex interplay among nervous, circulating, and cerebral immune cells, which can play dual roles influenced by their spatial location, context, and timing [34]. Despite the significance of these factors, many cytoprotective therapeutic approaches have overlooked them, instead focusing on time windows that do not align with the underlying pathophysiology. Typically, interventions target the hyper-acute stage to fit the time range of rtPA, or slightly later, without discriminating the different phases. Therefore, an increased awareness of the intricate spatiotemporal relationships between the nervous, vascular, and immune systems may prove essential for the design of novel therapeutic approaches to improve stroke prognosis.

In this context, our findings advocate that the neuropeptide cortistatin, which provides a crucial link between the CNS and the immune system, could be a promising therapeutic agent, positively influencing different aspects of stroke pathophysiology when administered at appropriate stages. Surprisingly, when cortistatin was given immediately after blood flow restoration, it worsened neuronal damage, motor performance, and neurological scores. The decreased microglial density with increased ameboid-like microglia in the core and the hyper-ramified microglial phenotype in the penumbra might be related to the potent immunomodulatory effects of cortistatin. The neuropeptide administered shortly after reperfusion may blunt the initial endogenous response of microglia that tries to limit the ischemic damage. Cortistatin may have prompted microglia to respond with an initially protective but later excessively reactive phenotype. Furthermore, the immediate cortistatin treatment appeared to hinder the initial astrocytic response, as evidenced by the disorganized astrocytic scar, preventing the formation of well-defined astrocytic borders, and potentially facilitating the expansion of neuronal damage.

Conversely, treatment with cortistatin at a slightly later stage (4h + 24h) did not reduce infarct volume or improve neurological outcomes. This treatment did not affect microglia density or morphotype in the core but induced a hyper-ramified microglia phenotype in the perilesional area. The early treatment with cortistatin enhanced the formation of the glial scar, possibly because it coincided with the initiation of the astrocytic response [37], modulating astrocytes differently than at the 0-hour time-point treatment. These findings underscore the critical importance of timing in therapeutic intervention, as even a few hours can have a strong effect on the response to the treatment. Indeed, late cortistatin administration (24h) appeared to lead to reduced neuronal lesions and improved outcomes.

The lack of protection observed with the immediate administration might seem contradictory to the beneficial effect shown at 24h and to the reported beneficial effects of cortistatin administration in other neurodegenerative/neuroinflammatory disorders [8,11,12]. However, the latter studies involved cortistatin injection at later stages of the disorder (*i.e.,* when the injury is well-established), while in our investigation the immediate administration coincided with the initial stages of injury. This suggests that the observed harmful effects on outcome are not likely due to cortistatin being neurotoxic but rather that when administered immediately or early after stroke, cortistatin interferes with the initial local neuroinflammatory response that is essential for containing the damage and removing debris and dead cells (*i.e.,* glial scar formation and microglia response, respectively), thereby leading to detrimental neurological outcome. The timing of cortistatin administration therefore appears vital to the subsequent progression of brain healing processes. Indeed, blocking or modifying the important early response leads to adverse outcomes in other disorders. For instance, Szalay *et al.* demonstrated that depletion of microglia resulted in a significant enhancement of lesion size, which was reversed by microglial repopulation [38]. These findings may also explain the lack of clinical success reported with some anti-inflammatory/immunomodulatory drugs administered at too early phases [3]. Collectively, these data underscore the fundamental role of the inflammatory response in the regenerative process after stroke and suggest that these mechanisms should not be completely halted. Instead, modulating the inflammatory response, rather than attempting to stop it entirely, seems to be a more effective approach.

Given these first results, we focused on a deeper investigation of the effect of cortistatin administration at the late stage. This approach, besides providing more flexibility in treatment application, also enables concurrent administration with other therapies such as rtPA. First, we confirmed the reduced striatal lesion size following late (24h) cortistatin treatment based on CV and MAP-2 immunodetection. Notably, MAP-2 provides a better insight into the first phases of neuronal death, even if the process is not yet concluded [39]. This indicated that cortistatin efficiently targeted neuronal death, protecting neuronal bodies, also evidenced by the increased expression of neuroprotective factors such as *BDNF* or *GDNF*. GDNF is important for the maintenance of dopaminergic neurons and their protection against injury [40], while BDNF protects neurons from ischemia-reperfusion-induced apoptosis [41]. The neuroprotective role of cortistatin conforms with previous research, where we and others demonstrated that treatment with the neuropeptide protected neurons from distinct insults *in vivo* and *in vitro* [12,42]. However, it is not clear yet the underlying mechanisms (*i.e.,* inhibition of apoptosis, neuronal degeneration, or oxidative damage). Moreover, we can hypothesize that cortistatin might also exert an indirect beneficial effect on neurons by regulating the responses of glial cells.

For decades, ischemic stroke research has primarily focused on rescuing neurons in the penumbra region to prevent the expansion of the infarct core. However, given that excitotoxicity, oxidative stress, neuroinflammation, or peripheral immune response contribute to secondary neuronal cell death [43] and that focusing solely on neuronal death is oversimplistic [44], this study aimed to explore the impact of cortistatin on the stroke-induced neuroinflammatory response. Microglia are the initial responders to ischemic damage, swiftly undergoing dynamic morphological and molecular changes in the infarct core and the peri-infarct zone [45]. Importantly, microglia undergo multiple alterations after injury (*e.g.,* degree and length of branching, branch configuration, shape and soma size, cell territory, receptor distribution, or cytoskeletal rearrangement), that go beyond the oversimplified paradigm of M1/M2 or the traditional morphological characterization categorizing microglia as either ramified (homeostatic) or reactive (short processes) [46]. Our findings showed an increased number of microglial cells in the ischemic core after late treatment with cortistatin. Moreover, elevated expression of markers related to phagocytosis (*i.e., CD68, Trem2*) was also found in the ipsilateral hemisphere of cortistatin-treated mice, consistent with the enhanced phagocytic phenotype and neuroprotective microglial function reported in early stages of stroke [47,48]. Since cortistatin significantly reduced brain lesions, these findings suggest an enhanced phagocytosis of dead neurons by the microglia from cortistatin-treated mice. Indeed, previous studies have demonstrated that endogenous cortistatin is essential for phagocytosis in physiological and demyelinating environments [49]. However, hyperreactive microglia over time can engulf live or stressed neurons, resulting in worsened neurological deficits [50], emphasizing the need to maintain a careful balance to ameliorate stroke prognosis. Microglia in the ischemic core exhibited reduced soma size and a larger number of junctions following cortistatin treatment, suggesting a shift to a more protective phenotype, diverging from the ameboid-like shape phenotype observed in control mice. While most microglia are engaged in engulfing dead neurons and secreting neurotrophic factors in the ischemic core, those in the peri-infarct area exhibit a different phenotypic polarization, influenced by surrounding factors [51,52]. Cortistatin treatment shifted peri-infarct microglia to a distinct reactive phenotype characterized by a reduced territory and soma but increased complexity. Moreover, elevated levels of *TNF-α*, *IL-10*, and neuroprotective factors such as *BDNF* or *GDNF* were detected. These results suggest a potential transformation to a distinct subtype of protective/reactive microglia, capable of phagocytosing cellular debris while aiding brain regeneration [53]. This is confirmed by previous findings in other neuroinflammatory models (*i.e.,* multiple sclerosis, Parkinson’s disease, and sepsis) [8,12,54], where cortistatin-modulated microglia markedly correlated with a better disease prognosis. Conversely, we observed a decreased expression of the scavenger receptor *CD163*, involved in the clearance of haemoglobin [55]. Interestingly, this factor is upregulated in patients with intracerebral haemorrhage and edema development [56], suggesting a potential reduction of these events following cortistatin treatment. Overall, the unique combination of phagocytic function, inflammatory cytokines, and neurotrophic/neuroprotective factors secreted by these cells may play a crucial role in promoting neuron survival and modulating other neuroinflammatory responses. Although many questions remain unanswered regarding the transitions between the different subsets of glial cells and the key molecular cues that drive these transitions [23], our study indicates that cortistatin may play an important role in this context. However, acknowledging the limitations of our study, we could not determine the exact molecular and functional morphotype of these cells in terms of spatial and temporal dynamics, since the gene expression analysis could not discriminate between different brain areas within the same hemisphere, different cell populations, or different cell subsets. Additionally, we cannot dismiss the fact that more than one cell type produces the same factor, further complicating the interpretation of our results. To address these limitations, further studies are needed to explore the exact mechanisms through which cortistatin-modulated microglia exert this immunomodulatory role.

Astrocytes also play essential roles in stroke, modulating the neuroinflammatory response, protecting neurons by releasing neuroprotective factors, forming barriers to limit damage, and promoting brain repair [57]. Although the molecular and cellular mechanisms that underlie the formation of the glial scar and astrocyte reactivity are not completely understood, they likely involve a complex and balanced interplay of molecular signals that can simultaneously boost phagocytosis and debris clearance, which would be essential for the protection and conservation of the still-healthy tissue [37,57,58]. Similar to microglia, an A1/A2 paradigm for astrocyte morphology and function has been proposed [59,60]. However, the spatiotemporal heterogeneity of reactive astrocytes has demonstrated that this classification is too simple, so additional parameters should be considered [61]. Our study revealed that cortistatin-treated mice exhibited a well-organized and thicker glial scar than control mice. This was accompanied by an increased expression of neurotrophins, which have been described to be involved in synaptic plasticity and neuronal growth regulation that tend to decrease in pathological CNS processes [62]. Thus, cortistatin promotes a distinct reactive phenotype that supports tissue repair and neuroprotection. An effective glial scar is critical as defective scar formation has been associated with larger lesions, demyelination, and reduced neuronal survival and functional recovery [37,58,63]. However, an excessive glial scar can be detrimental by exacerbating inflammation and interfering with synaptic and axonal growth [37]. The observed localization and cortistatin-dependent timing of astrocyte activation in our study indicate a controlled and neuroprotective response to limit damage following cortistatin treatment.

In addition, BBB integrity is needed for better stroke outcomes [64]. The traditional classification of BBB changes during stroke progression is restricted to “opening” or “closing” (*i.e.,* "bad" or " good”) making this dichotomy outdated [65]. Instead, BBB permeability follows a sequence across different phases (*i.e.,* hyperacute, acute, subacute, and chronic phases), each of them characterized by distinct biological events [65]. Our study specifically focused on the acute phase in which BBB permeability peaks, facilitating neuroinflammation progression and exacerbating injury [66]. Our findings demonstrated that cortistatin treatment helped preserve BBB integrity by reducing endothelial dysfunction and maintaining tight junctions, as evidenced by the decreased IgG leakage and reduced number of ZO-1 gaps. These results align with previous studies showing that cortistatin protects the BBB in models of mild neuroinflammation and *in vitr*o models of BBB disruption [13]. Interestingly, the lack of changes in claudin-1 and claudin-5 gene expression suggests that TJ alterations may occur at the protein localization level rather than at the gene expression level [67–69]. Indeed, cortistatin has demonstrated a role in modulating protein levels, cellular redistribution, and functional organization of BBB proteins such as ZO-1, claudin-1, claudin-5, and VE-cadherin without significantly affecting their transcript levels [13].

Finally, it is important to acknowledge that stroke is a complex, organism-wide process that not only targets the brain but also peripheral tissues. Shortly after the interruption of the CBF and subsequent hypoxia, ECs express selectins in response to oxidative damage, quickly engaging circulating innate immune cells, and triggering a rapid pro-inflammatory response in the periphery [2]. The breakdown of various components of the BBB enables the extravasation of these cells, with different cadences and at variable times. In general, a well-regulated inflammatory response is inherently self-limiting, facilitating the repair, turnover, and tissue adaptation, and prioritizing processes that restore homeostasis over those that are harmful [70,71]. The immunomodulatory effects of cortistatin were evident as it reduced immune cell infiltration into the brain and modulated systemic immune responses, consistent with its effects observed in other neuroinflammatory models [8,13]. Specifically, cortistatin treatment modulated the expression of typically pro-inflammatory cytokines (*e.g.,* IL-6, TNF-α, IL-12) in the brain. These cytokines exert biphasic or even polyphasic roles in ischemic stroke, and depending on the timing, spatial location, and severity of the injury, they can play vital roles in repair processes or have detrimental effects. For example, it has been shown that the deletion of IL-6 in MCAO mice does not confer protection against cerebral ischemia [70]. Indeed, IL-6 serves as a neurotrophic factor, improving functional outcomes, reducing stroke volume in mice [72], and promoting post-stroke angiogenesis [73]. Similarly, numerous studies have reported a neuroprotective role of TNF-α in reducing the production of nitric oxide and free radicals, maintaining neuronal calcium homeostasis, and inducing the synthesis of neurotrophic factors [74]. Hence, these data, along with the increased levels of anti-inflammatory factors (*i.e.,* IL-10), the reduced number of infiltrating cells, and the decreased neuronal damage suggest that the elevated inflammatory factors observed after the treatment with the neuropeptide might be crucial for developing an appropriate response to the acute damage and subsequent resolution rather than playing a detrimental role. This underscores the complexity of the immune response in stroke and, again, the importance of the timing of therapeutic intervention.

This initial activation of systemic immunity is quickly succeeded by a prolonged state of immunodepression, which enhances the susceptibility to infection or other pathologies [36]. Depression of the immune system not only occurs in patients with ischemic stroke but also in other acute CNS injuries such as cerebral haemorrhage, traumatic brain injury, or spinal cord trauma (reviewed in [36]). In this sense, the observed potent immunomodulatory role of cortistatin, especially regarding spleen function and cytokine recovery to homeostatic levels, is consistent with other studies [8,75,76]. Interestingly, we found elevated levels of MCP-1 after cortistatin treatment. Although this factor is systemically upregulated in stroke patients and is known to promote leukocyte chemotaxis, some reports also demonstrate that MCP-1 is crucial for recruiting neural stem cells after stroke, which is necessary for better neurological outcomes [77]. Along these lines, we also found that cortistatin reduced the elevated levels of *Arg1*, which is consistently upregulated in stroke patients [32], and known to induce lymphopenia in a murine model of stroke [78]. Indeed, Arg1 inhibition ameliorates stroke outcomes [79] and it is already been used in clinical trials in patients with coronary artery disease who have suffered from ischemia-reperfusion [80].

In summary, with timely administration, the neuroimmune-modulatory role of cortistatin in stroke extends beyond its direct neuroprotective effects on brain damage, glial responses, and BBB modulation, to include the regulation of systemic immune responses and to positively affect detrimental behavioural and functional outcomes. This multifaceted mechanism of action offers several advantages over therapies targeting single aspects of stroke pathophysiology. Furthermore, the intriguing possibility that this neuropeptide has a role in the main players in this condition (such as neurons, astrocytes, microglia, BECs, pericytes, or immune cells) could confer an advantage over other potential therapies. Notably, the physiological nature—and thus intrinsically non-toxic properties—of this neuropeptide provide assurances of its potential use. In fact, it has been successfully used at clinical levels, both systemically and locally, demonstrating its therapeutic efficacy in patients with acromegaly, prolactinoma [81], and Cushing’s disease [82]. Being beyond the scope of this work, further research should consider sex as an independent variable that must be addressed to properly interpret the beneficial action of cortistatin. Future studies should also focus on extending the investigation of the spatiotemporal dynamics of cortistatin action, the molecular pathways involved, and its effects on other cell types within the CNS and the peripheral immune system.

## CONCLUSION

Our data underscore the importance of comprehending the spatio-temporal dynamics of ischemic stroke when devising therapeutic interventions to optimize stroke outcomes. The findings indicating that a too-potent anti-inflammatory effect of the neuropeptide is deleterious when given immediately after stroke, emphasize the need to shift therapeutic goals from halting specific processes to modulating and balancing them, thereby mitigating possible chronic adverse responses. We showed that cortistatin administered at later stages appears to be a very promising candidate drug to enhance recovery after stroke, alone or in combination with other treatments, mainly targeting the outcome of patients that underwent recanalization or spontaneous reperfusion. Our study further highlighted the multifaceted roles of cortistatin in stroke pathophysiology, emphasizing its potential as a therapeutic agent. Its capacity to simultaneously act on various pathological features, such as neuronal damage, glial over-reaction, astrocytic scar formation, and BBB breakdown with an overall amelioration of the neurological outcome, offers significant advantage over interventions targeting single aspects of this condition. Moreover, cortistatin stands as a key factor regulating the complex interplay between the CNS and the immune system, modulating exacerbated molecular and cellular responses from both systems that may affect the brain’s recovery processes. Finally, the valuable insights gained from this research have the potential to advance the development of novel therapeutic approaches, not only targeting stroke pathology but also other CNS disorders linked to glial over-reactivity, BBB disruption, or dysfunctional CNS-immune system interactions. Future research should continue to explore the complex interplay between this neuropeptide, its receptors, and the multitarget effects in stroke to develop more effective therapeutic strategies.

## ACKNOWLEDGEMENTS

The authors would like to thank the Animal Care Unit and Microscopy Service Unit staff from IPBLN for the technical support.

## CONTRIBUTIONS

E.G-R conceived, designed and supervised the whole study. J.C-G and E.G-R outlined the experiments and analysed all the data. J.C-G, L.B, P.V-R, I.S-M, and I.F-L performed the experiments. J.C-G, L.B, M.P, P.H-C, L.H, and E.G-R interpreted the data, and discussed the results. J.C-G and E.G-R wrote and drafted the manuscript. J.C-G, E.G-R, L.B, M.P, and L.H revised the manuscript. All authors have read and agreed to the published version of the manuscript.

## ETHICS DECLARATION

### Ethical approval

All procedures were approved by the Animal Care and Use Board and the Ethical Committee of the Spanish National Research Council (Spain), and the Vaud Cantonal Veterinary Office (Switzerland). All experimental protocols adhered to the guidelines specified in Directive 2010/63/EU of the European Parliament and the corresponding Swiss regulations regarding animal protection for scientific purposes.

## DATA AVAILABILITY

All data used or analysed will be made available upon reasonable request to the corresponding author.

## ABBREVIATIONS

Arg1: arginase 1
BBB: blood-brain barrier
BDNF: brain-derived neurotrophic factor
BECs: brain endothelial cells
BL: baseline
CBF: cerebral blood flow
Cl: claudin
CD: cluster of differentiation
CL: contralateral
CNS: central nervous system
CST: cortistatin
CV: cresyl violet
DAMPs: danger-associated molecular pattern molecules
GDNF: glial cell-derived neurotrophic factor
GFAP: glial fibrillary acidic protein
Iba1: ionized calcium-binding adaptor molecule 1
IgG: immunoglobulin G
IL: Interleukin
MAP-2: microtubule-associated protein 2
MBP: myelin basic protein
MCAO: middle cerebral artery occlusion
MCP-1: monocyte chemoattractant protein-1
PFA: paraformaldehyde
Rplp0: ribosomal protein lateral stalk subunit P0
rtPA: recombinant tissue plasminogen activator
TJ: tight-junctions
TNF: tumour necrosis factor-α
Trem2: triggering receptor expressed on myeloid cells 2
ZO-1: zonula occludens 1

